# Assessment of adult structural plasticity in *Drosophila* neurons

**DOI:** 10.64898/2026.03.11.711108

**Authors:** Micaela Rodriguez-Caron, Francisco J. Tassara, Juan I. Ispizua, Christian M. Carpio-Romero, M. Fernanda Ceriani

## Abstract

Unraveling how adult neurons reshape their architecture is key to understanding post-developmental plasticity. *Drosophila* clock neurons, which remodel their terminals on a daily basis, offer a unique model to examine the mechanisms underlying structural plasticity. In this study, we examine the impact of the experimental design on the remodeling process. We established a simple fixation protocol that preserves tissue integrity and prevents its deformation while enabling the fixation of a larger number of individuals within the appropriate time window. We show that intrinsic (i.e., targeting fluorescent reporters to the membrane) or extrinsic (i.e., temperature) variables may influence this dynamic process. Examining *ex vivo* preparations, we found that the s-LNv terminals display numerous thin filopodia extending from their synaptic boutons. However, these fine membrane protrusions are lost upon fixation, as they could only be accurately visualized *ex vivo*. Finally, we present *MorphoScope*, a Python-based interface that eliminates observer bias in complexity measurements. Altogether, we present a powerful and robust model to investigate the principles of adult neuronal plasticity, with implications extending beyond circadian biology.

## Introduction

Neuronal plasticity, the ability of the nervous system to modify its structure and function, underlies fundamental processes such as learning, memory, and adaptation to changing environments. While mostly characterized during development and early critical periods, plasticity also persists in the adult brain, though often in more subtle and spatially restricted form. This phenomenon is conserved across cell types and species: olfactory systems adjust according to age, sensory experience, and internal state (Anton & Rössler, 2021); astrocytes restructure their processes to regulate synaptic remodeling and behavior (Lawal et al., 2022); and rhythmic influences, such as circadian or homeostatic drives, induce dynamic changes in neuronal morphology. In *Drosophila*, the projections of dopaminergic neurons adjust according to nutritional state, and the size and shape of their optic lobe neurons change on a daily basis (Pyza & Meinertzhagen, 1999; Liu et al., 2017). In zebrafish, hypocretin neurons modulate the number of synapses according to circadian time and sleep pressure (Appelbaum et al., 2010). Moreover, the mammalian suprachiasmatic nucleus undergoes daily remodeling of neuronal structure, as well as glial coverage of neuronal terminals (Becquet et al., 2008; Neitz et al., 2024).

Within this spectrum, structural plasticity refers to the dynamic changes in the morphology of neurons, including the growth, retraction, and remodeling of dendritic and axonal processes. These changes are central not only to normal brain function but also to aging and the pathophysiology of psychiatric and neurological disorders (Barnes, 2001; Bernardinelli et al., 2014). Therefore, it is essential to identify tractable models in which to study structural plasticity in the adult brain, and *Drosophila melanogaster* circadian neurons have emerged as a powerful one. The small lateral ventral neurons (s-LNvs), that express the neuropeptide pigment-dispersing factor (PDF), display daily structural changes in their axonal projections (Fernandez et al., 2008). This remodeling is regulated by the cell-autonomous circadian clock, activity-dependent mechanisms, and even PDF signaling, but it is also shaped by glia and communication with additional circadian clusters (Depetris-Chauvin et al., 2011; Sivachenko et al., 2013; Petsakou et al., 2015; Gunawardhana & Hardin, 2017; Herrero et al., 2017, 2020; Polcowñuk et al., 2021; Damulewicz et al., 2022; Vaughen et al., 2022; Lymer et al., 2024). Although distinct scenarios might require some system-specific components, structural plasticity likely follows general guiding principles and recruits common mechanisms. Hence, daily restructuring of the s-LNv axonal arbor provides a uniquely dynamic, experimentally and temporally accessible system for dissecting the principles of adult structural plasticity. However, after years of working with this phenomenon, we noticed that its analysis is constrained by unrecognized sources of variability. This highlights the need for more robust methodological standards.

Here, we examine different possible sources of variability in s-LNv structural plasticity. We optimized fixation and mounting protocols to preserve brain and neuronal structure and ensure consistent access for image acquisition. We demonstrate that neuronal labeling strategies have a significant impact on remodeling, with combined membrane and cytoplasmic markers proving to be particularly disruptive. We explore the impact of routinely used temperatures and light intensities, finding that light has no significant effect on the rhythm amplitude, whereas temperature exerts a subtle yet global influence on structural complexity. Additionally, we introduce *MorphoScope*, a Python-based interface to facilitate structural complexity quantification while reducing observer biases. Together, these advances provide a framework for using s-LNvs as a model to investigate mechanisms underlying structural plasticity in the adult brain.

## Results

### Tissue fixation and mounting media impact overall brain morphology

The description of complex axonal or dendritic arbors is usually performed in fixed tissue. Fixation is necessary to prevent autolysis and degradation, thereby preserving tissue components for subsequent immunolabeling and fluorescence microscopy. Among the numerous available fixatives, paraformaldehyde (PFA) is one of the most widely used. Although PFA concentrations and general procedures are relatively standardized, subtle variations in how the tissue is exposed to the fixative -whether it is first dissected, the incubation length or temperature- can affect tissue integrity and labeling outcomes, consequently affecting the quantification of plasticity and reproducibility of the results.

Most commonly used immunostaining protocols for adult *Drosophila* brains involve dissecting unfixed brains prior to fixation (Wu & Luo, 2006; Flylight project, 2020). This approach has several drawbacks. When dissections are performed within a limited time window (i.e., at a given circadian time), this restricts the number of samples that can be processed, making it difficult to integrate multiple genotypes or conditions for comparison within the experiment. Moreover, unfixed brains are fragile and prone to mechanical damage or distortion during dissection. To overcome these limitations, we compared the impact on brain morphology of three fixation strategies normally used in our laboratory that do not require dissection as the first step. Anesthetized flies were decapitated and their heads fixed in 4% PFA at room temperature (RT) for 50 minutes or kept in the fixative on ice for 60 minutes. Alternatively, whole flies were briefly de-waxed in 96% ethanol before fixation in 4% PFA at RT for 60 minutes. The first approach produced brains with well-preserved overall morphology and reliable immunostaining, characterized by rounded varicosities and sharply defined thin processes (**Figure 1A**). On the other hand, both ice fixation and the ethanol pre-treatment led to a high incidence of deformed brains and poor-quality immunofluorescence (**Figures 1B and 1C**), presenting abnormally stretched terminals and diffuse processes, which likely reflects suboptimal or incomplete fixation.

**Figure 1.**
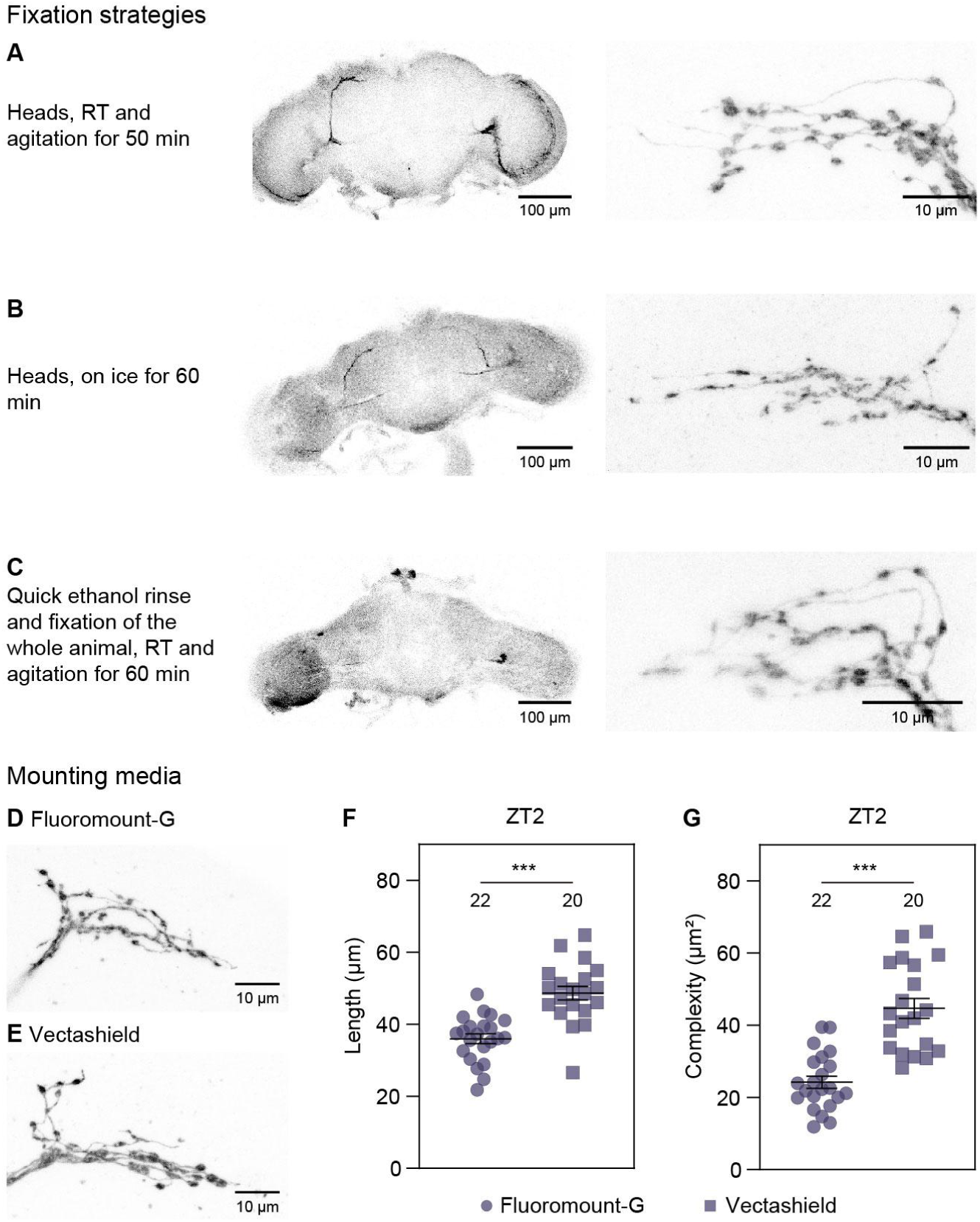
Tissue fixation and mounting media impact overall brain morphology. **(A-C)** Representative images of the general brain and terminal morphology from *Pdf-*GAL4 > UAS-CD8::GFP, *Pdf*-RFP animals. Heads were fixed in 4% PFA at RT with gentle agitation **(A)**, fixed on ice without agitation **(B)** or de-waxed whole animals were fixed at RT with gentle agitation **(C)**. RFP fluorescence is shown in the right panels. **(D, E)** Representative terminal morphology from *Pdf*-RFP brains dissected at ZT2 and mounted in either Fluoromount-G **(D)** or Vectashield **(E)**. **(F, G)** Quantification of terminal length **(F)** and complexity **(G)** in samples mounted in Fluoromount-G and Vectashield. Each dot represents the terminals of a single hemibrain per fly. The number of hemibrains (n) analyzed in each case is indicated above the symbols. Error bars represent the SEM. Asterisks indicate statistically significant differences: * p < 0.05, ** p < 0.01, *** p < 0.001. Full brain images are single planes, while terminal images are Z-projections of Z-stacks.

In addition to fixation, we found that the choice of mounting media acts as an additional source of morphological variability. To evaluate this effect, we quantified the length and structural complexity of terminals in brains dissected at ZT2, comparing two common commercial media: Vectashield and Fluoromount-G. Along this work, structural complexity (XY spread) was quantified using *MorphoScope*, a custom Python interface built upon a previously published script (Petsakou et al., 2015) that was modified to include additional functionalities. This tool was designed for intuitive use and real-time visualization and, contrary to the widely accepted Sholl method, it is not strongly influenced by observer bias (see **Supplementary Figure 1** and **Materials and Methods**).

We found that terminals mounted in Vectashield displayed significantly greater length and complexity than those in Fluoromount-G (**Figures 1D-1G**). Although there is no intrinsic reason to prefer one medium to another, Vectashield appears to provide better resolution of the finest neuronal processes, making it a suitable choice for evaluating delicate structures. However, its fluid nature might enable tissue flattening over time. In contrast, Fluoromount-G solidifies, which likely constrains and maintains the tissue structure more effectively. Although hard-setting variants of Vectashield could be used as an alternative, the mounting medium selected must remain constant throughout experiments to avoid artifactual variability. Finally, re-imaging the same samples after two weeks revealed a slight reduction in the XY spread (**Supplementary Figure 2**). While this difference did not reach statistical significance, it underscores the importance of minimizing the time elapsed between mounting and acquisition.

Overall, these findings highlight the critical role of both fixation conditions and mounting media in achieving consistent brain morphology and reliable immunofluorescence results. The standardization of these steps should result in a reliable protocol that allows the processing of larger sample numbers within constrained time windows, thus minimizing both tissue damage and methodological variability.

### Fluorescent reporter choice affects neuronal remodeling

Multiple markers have been used to describe the extent of circadian structural remodeling, ranging from direct neuropeptide immunolabeling to membrane-targeted versions of commonly used fluorescent reporters (Fernandez et al., 2008; Herrero et al., 2020). We compared three distinct strategies to label the neuronal architecture of the s-LNv terminals in order to evaluate their suitability as inputs for structural complexity quantification. These included GAL4-mediated expression of a membrane-tethered GFP, expression of a cytoplasmic RFP under the *Pdf* promoter, and immunolabeling of the endogenous PDF neuropeptide. Flies were evaluated at two key time points, two hours after lights on (*Zeitgeber Time* 2, ZT2) and two hours after lights off (ZT14), using either one or both reporters in combination with PDF immunostaining to allow direct comparison of the different labeling strategies.

First, we assessed complexity through quantification of the XY spread of the s-LNv terminals labeled with cytoplasmic RFP and PDF immunoreactivity. Both markers reliably reported structural differences between morning and night across hemispheres (**Figure 2A**). No differences were observed when PDF images were acquired employing the optimal confocal settings per sample (as shown in the left hemispheres) or when acquired using the optimal settings for the ZT2 samples (as customary when comparing PDF intensity, right hemispheres, see **Materials and Methods**). In parallel, we assessed structural plasticity through the expression of a membrane reporter along with the endogenous PDF signal in *Pdf*-GAL4 > UAS-CD8::GFP flies. Surprisingly, under these conditions, the analysis of the GFP signal retrieved negligible structural changes between early morning and early night (**Figure 2B**). However, the PDF signal still revealed significant differences in complexity between the two time points, perhaps as a result of its cyclical nature (**Figure 2A**). To address if the inability to detect structural changes is the result of abnormal reporter expression or a more general effect on neuronal physiology we combined the membrane and cytoplasmic markers in *Pdf-*GAL4 > UAS-CD8::GFP, *Pdf*-RFP flies. Under these conditions, no changes in the degree of arborization were observed regardless of the signal employed during the quantification (**Figure 2C**). The comparison of structural plasticity through RFP quantification in the *Pdf*-RFP (**Figure 2A**) and *Pdf-*GAL4 > UAS-CD8::GFP, *Pdf*-RFP (**Figure 2C**) strains shows differences at ZT2, where animals expressing both the membrane and cytoplasmic reporters exhibited terminals with reduced complexity (see **Supplementary Table 1**). Altogether, these results suggest that membrane-targeted reporter expression overloads the structure and in turn disrupts normal daily remodeling without obliterating PDF cycling (not shown).

**Figure 2.**
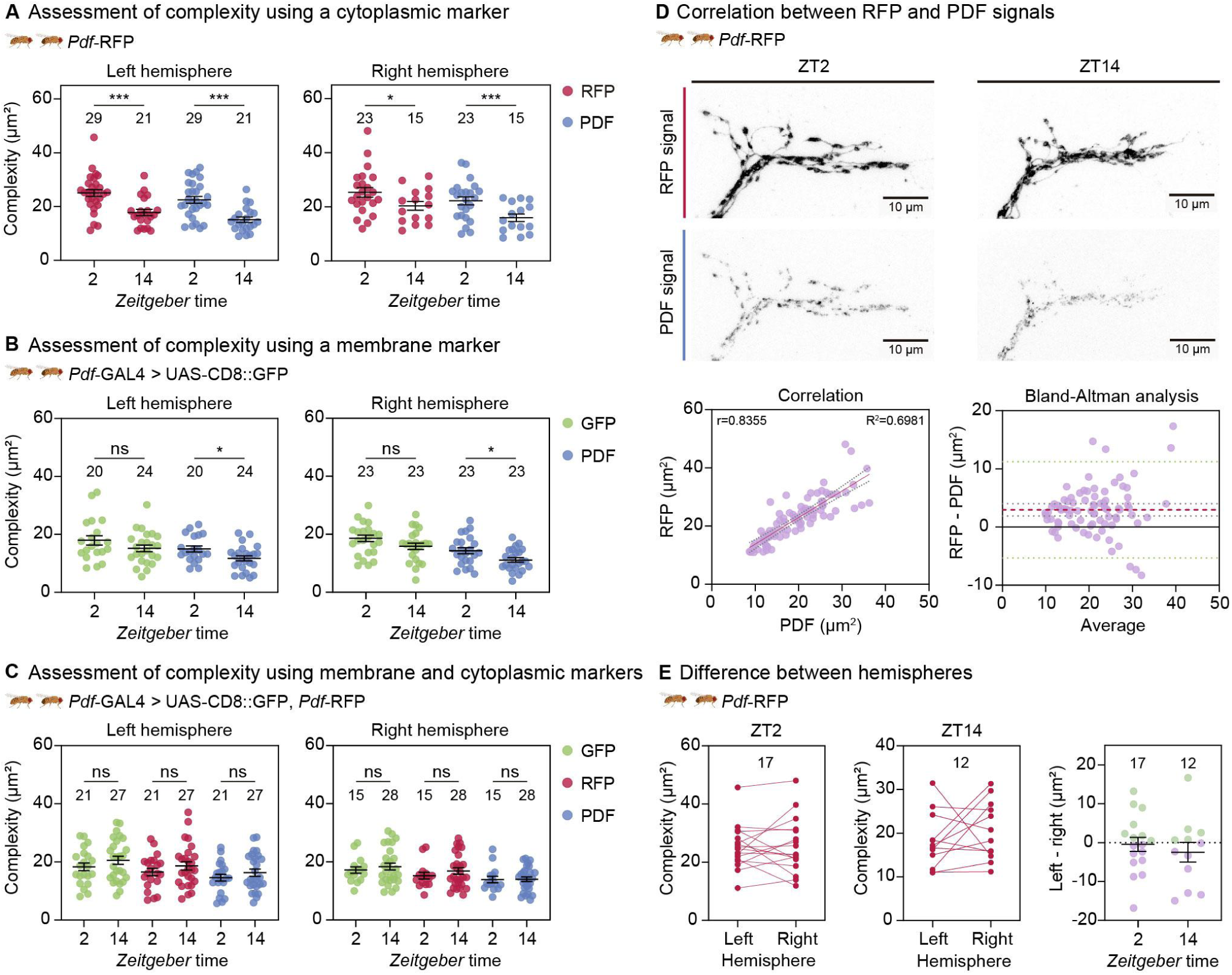
Fluorescent reporter choice affects neuronal remodeling. **(A-C)** Graphs show structural complexity (XY spread) at two key time points, ZT2 and ZT14, in both brain hemispheres, using three different reporters as the input for quantification: membrane GFP, cytoplasmic RFP, and endogenous PDF. The PDF signal was compared to RFP **(A)**, GFP **(B)** and the combined expression **(C)**. Each dot represents the terminals of a single hemibrain per fly. **(D)** Comparison of cytoplasmic and PDF signals. Top panels show representative images of the same terminal described by RFP (top) or PDF (bottom) at ZT2 and 14. Bottom panels show the correlation analysis of the two signals (left) and Bland-Altman analysis to compare agreement between the two reporters (right). The difference between the complexity measured with the cytoplasmic reporter and PDF is plotted against their average. The red dashed line represents the bias, while the green lines represent the standard deviation of that bias (that is, the limits of agreement). The grey dotted lines represent the 95% of the confidence interval of the mean. In both cases, each dot represents one hemibrain. **(E)** Assessment of the complexity in the left and right hemispheres at ZT2 (left panel) and ZT14 (middle panel) in *Pdf*-RFP animals. Lines connect the two hemispheres of an individual brain. The right panel shows the difference in complexity between the two hemispheres (left–right) at either time point, with each dot representing one brain, each from a different fly. The number of brains or hemibrains (n) analyzed in each case is indicated above the symbols. Error bars represent the SEM. Asterisks indicate statistically significant differences: * p < 0.05, ** p < 0.01, *** p < 0.001, ns = not significant.

Although the PDF signal appears to correctly reflect the changes in structure complexity, we further assessed its accuracy by direct comparison to the cytoplasmic RFP. As shown in **Figure 2D** (upper panel), the RFP signal provides a more detailed description of the overall structure both at ZT2 and ZT14, whereas the punctated PDF signal fails to capture thinner processes. We then performed a correlation analysis between the complexity scores obtained from both signals within each hemibrain included in **Figure 2A**. As expected, this analysis revealed a strong positive relationship (r = 0.8355, left bottom panel, **Figure 2D**). However, while correlation assesses the strength of the relationship between two variables, a high correlation does not necessarily indicate that the two methods are in agreement. To perform a more thorough evaluation of the coherence between them, a Bland-Altman analysis was conducted (Giavarina, 2015). In this analysis, the difference between the two measurements for the same brain is plotted against their average. As shown in the bottom right panel of **Figure 2D**, the line of equality (y = 0) lies outside the range defined by the confidence intervals of the mean difference (grey dotted and red dashed lines, respectively). This suggests there is a systematic difference between the two methods. Specifically, there is a small -yet significant- positive bias, with slightly higher RFP values than those of PDF, which was also evident in the correlation analysis. While most values fall within the 95% limits of agreement (green dotted lines), this systematic difference indicates that PDF captures the overall structural change but with reduced accuracy compared to the cytoplasmic reporter.

Finally, the direct comparison of the complexity of the terminals between the left and right hemispheres revealed no consistent asymmetry either at ZT2 or ZT14 in *Pdf*-RFP flies (**Figure 2E**). Paired analyses showed variability across individuals, but the mean left-right difference was not significantly different from zero at either time point (**Figure 2E**, right panel), indicating that structural remodeling occurs similarly in both hemispheres.

### Exploring the effect of environmental conditions

It has been proposed that circadian structural plasticity integrates environmental cues such as light and temperature (Petsakou et al., 2015; Fernandez et al., 2020). Here, we examined whether light intensity and temperature conditions commonly used in the laboratory directly influence this remodeling process. *Pdf*-RFP-expressing flies were reared at 22°C under a low-light LD 12:12 regime (70-280 lux). Then, they were transferred to either low (70-280 lux) or high (1500-1600 lux) light intensity either at 22°C, 25°C, or 28°C. Single experiments including 8-14 brains per condition (**Supplementary Figure 3A and 3B**) show rather inconsistent results. This lack of reproducibility prompted us to estimate the minimal sample size required to achieve sufficient statistical power and avoid concluding that structural remodeling in sLNvs terminals is affected when it is not. Using the pooled *Pdf*-RFP data collected at 25°C (**Figures 2 and 3**), a power analysis indicated that a minimum sample size of 16 brains per group is necessary to achieve 80% statistical power (p < 0.05, **Supplementary Figure 3**) for two-sample comparisons under these specific conditions. Concurrently, the magnitude of the effect was determined using the standardized parameter known as Cohen’s d. For 16 animals per time point, the calculated Cohen’s d value approaches 1, a value that indicates a substantial effect size (Cohen, 1988). When the two independent experiments were pooled, ANOVA revealed no significant interaction between factors (**Supplementary Table 1**). Light intensity had no significant effect (p = 0.4731), and hence data from both light conditions was combined. The final model included time of day (p < 0.001) and temperature (p = 0.006) as significant factors, indicating that terminals are on average 6.24 µm² more complex at ZT2 than at ZT14. The effect of temperature, although subtle, suggests slightly higher complexity at 22°C compared to 25°C (by 2.87 µm²) and 28°C (by 2.32 µm²), which represents a global change in average complexity rather than an alteration of the remodeling between time points.

**Figure 3.**
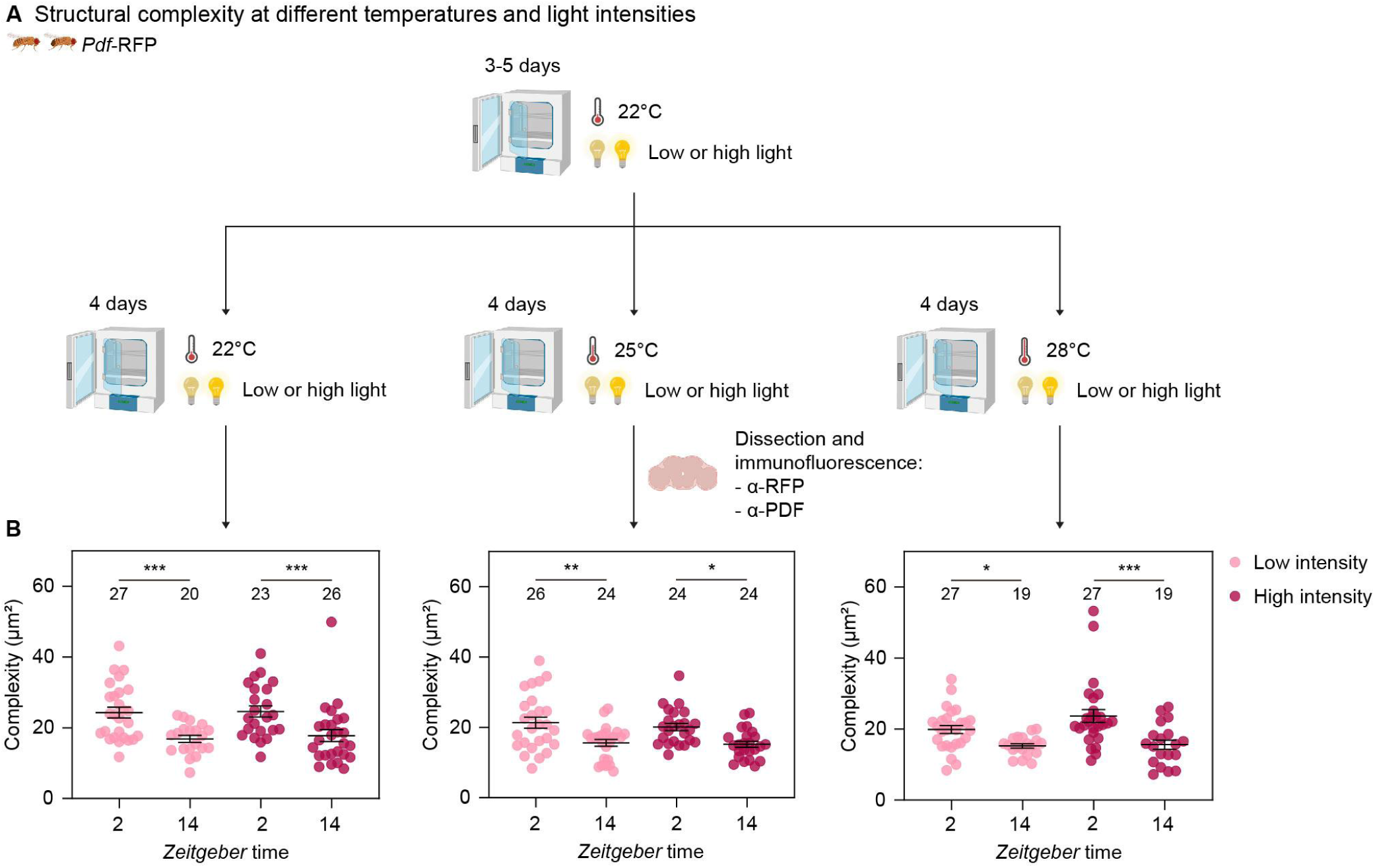
Exploring the effect of environmental conditions. **(A)** *Pdf*-RFP flies were maintained at 22°C under low light intensity for 3-5 days, and then either kept under the same condition or transferred for 4 days to environments with low or high light intensity at 22°C, 25°C, or 28°C. Brains were subsequently dissected and processed for immunofluorescence. Schematic diagram created in Biorender.com. **(B)** Data from two independent experiments were pooled and analyzed together. As light intensity was not a significant factor in the ANOVA (p = 0.473), both light conditions were combined; graphs still display data separated by time point, light intensity and temperature in order to visualize the different conditions. Each dot represents the terminals of a single hemibrain from one fly. Sample sizes (n) are indicated above each group. Error bars represent the SEM. Asterisks indicate statistically significant differences: * p < 0.05, ** p < 0.01, *** p < 0.001, ns = not significant.

Together, these results indicate that neither light intensity nor temperature affect the remodeling process itself, although temperature appears to exert a modest global effect on overall structural complexity, with animals kept at lower temperatures exhibiting more complex terminals.

### Newly formed protrusions are particularly sensitive to manipulation

Filopodia are thin and highly dynamic membrane protrusions that are formed during development and are required to explore and connect to the environment (Blake & Gallop, 2023). Filopodia mainly contain bundled actin filaments, whose delicate structure makes them highly susceptible to damage or alteration during standard tissue processing. They are likely the origin of the highly arborized structures observed in the s-LNv neurons in the early morning (Ispizua et al., 2025). Here, we compared the s-LNv terminals of animals expressing both CD8::GFP and *Pdf*-RFP in an *ex vivo* preparation and after undergoing a full immunolabeling procedure. In the *ex vivo* brains (**Figure 4A**), we observed thin protrusions labeled by the membrane marker (left panel) but not filled with cytoplasmic RFP (middle panel, compare varicosities in insets 1-3). This suggests that the former more accurately reports the overall structure. However, the membrane signal is strongly affected by standard immunolabeling protocols, resulting in the loss of a substantial number of filopodia in comparison to the *ex vivo* preparation (left panels, **Figure 4A and 4B**). This inability to preserve these delicate structures is likely due to fixation, as well as to the long incubation steps and washes with detergent-containing buffers. Conversely, the cytoplasmic reporter identifies similar structures in both preparations, consistent with the notion that nascent membrane protrusions are particularly fragile (middle panels, **Figure 4**). Taken together, these observations suggest that, to reliably study filopodia dynamics and plasticity, a membrane-targeted marker should be used in preparations without immunostaining.

**Figure 4.**
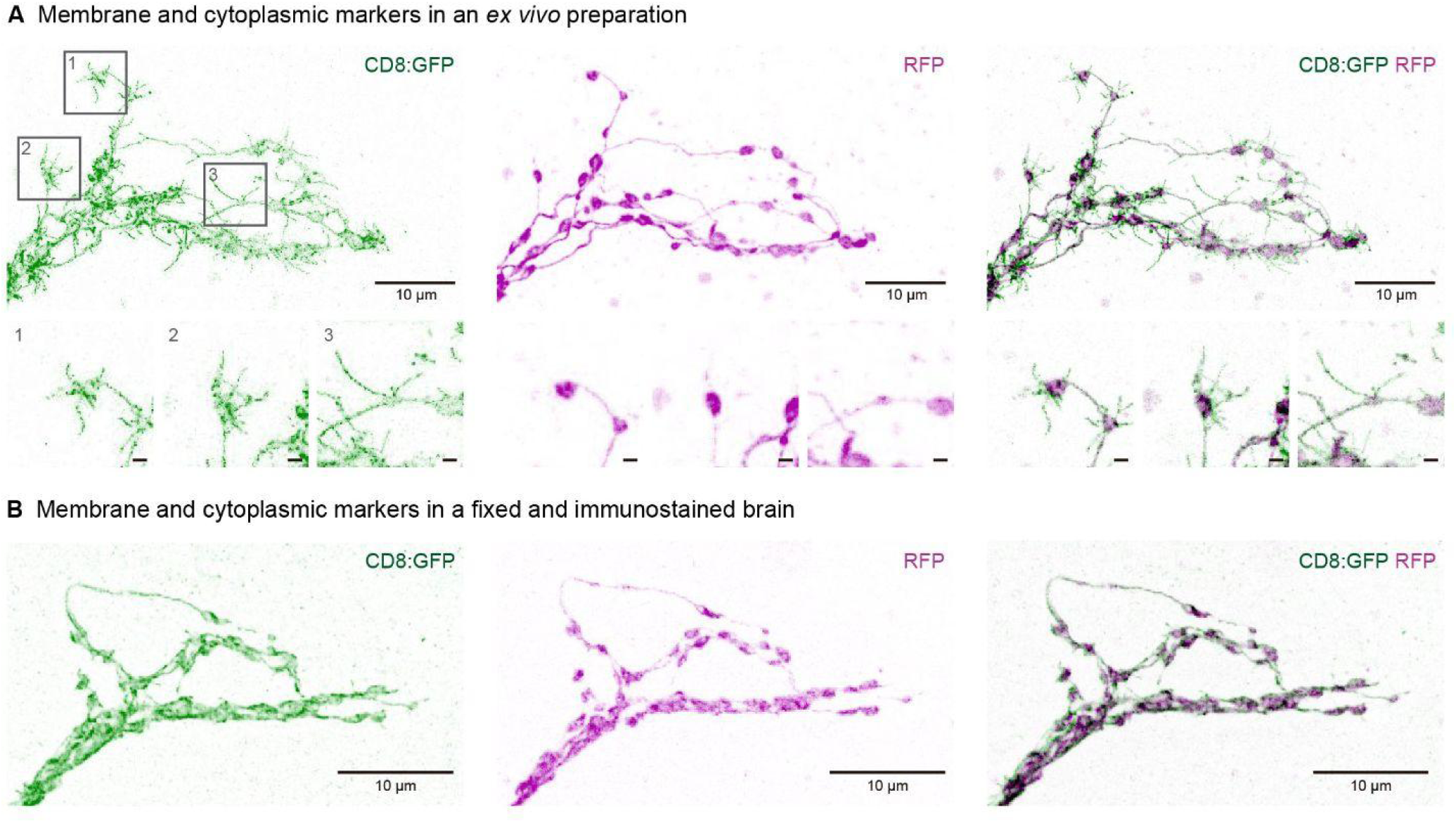
Newly formed protrusions are particularly sensitive to manipulation. **(A and B)** Comparison of membrane and cytoplasmic markers in an *ex vivo* preparation and after the optimal immunofluorescence procedure, respectively. From left to right: CD8::GFP (membrane marker), RFP (cytoplasmic marker), and the merged image. Insets **1, 2 and 3** show individual varicosities. Scale bars = 1 μm.

## Discussion

Circadian structural plasticity in *Drosophila* s-LNvs refers to the rhythmic remodeling of their axonal terminals throughout the day-night cycle (Fernandez et al., 2008). It results from the daily addition and pruning of varicosities, functional units containing all the elements required for effective communication: mitochondria, dense core vesicles, and synapses (Ispizua et al., 2025). Thus, remodeling involves dynamic changes in synaptic organization and neuronal complexity that depend on a functional circadian clock, cell-autonomous activity-dependent and independent mechanisms, and a continuous communication with neighbouring clock neurons and glia (Krzeptowski et al., 2018). These oscillations were proposed to adjust connectivity of the circadian network daily, making the s-LNvs a powerful model to study the mechanisms of adult neuronal plasticity in an accessible time scale. However, as with any phenomenon whose assessment relies on indirect variables, its evaluation is constrained by technical variability. Here, we identify some of the methodological sources of this variability in the *Drosophila* circadian circuit and provide a more detailed framework for studying the remodeling of the s-LNvs. By systematically comparing different parameters we show that experimental choices not only might affect reproducibility but also could interfere with the remodeling process itself. We also introduce *MorphoScope*, an interface that standardizes morphological analysis and minimizes observer bias often associated with broadly adopted strategies to quantify the degree of arborization such as Sholl analysis and related Fiji plug-ins (Sholl, 1953; Schindelin et al., 2012; Ferreira et al., 2014). Together, we are persuaded these advances will strengthen the adoption of the s-LNvs as a robust model to explore the principles of adult structural plasticity.

Structural plasticity depends on the coordination among membrane dynamics, cytoskeletal remodeling and intracellular signaling. Our study reveals that expression of the membrane-tethered reporter CD8::GFP could interfere with the daily remodeling of s-LNv terminals, whereas the cytoplasmic reporter preserves it. Different mechanisms could account for this effect. One possibility is the excessive accumulation of a membrane protein that increases macromolecular crowding, thereby altering the biophysical properties and dynamics of the plasma membrane. High protein density modulates intermolecular interactions, promoting nonspecific associations and clustering, or even changes in lipid packaging, which collectively could result in altered molecular mobility and diffusion properties, ultimately modifying membrane morphology and as such influencing remodeling (Frick et al., 2007; Löwe et al., 2020).

Given the relevance of activity-dependent mechanisms, particularly during the increase in terminal complexity characteristic of the early day (Depetris-Chauvin et al., 2011; Sivachenko et al., 2013; Lymer et al., 2024), another possibility is that excess of the membrane-tethered reporter would trigger changes in the membrane electrical properties. Although direct measurements in cultured cells with overexpressed proteins show that membrane capacitance is relatively robust (Gentet et al., 2000), *in vivo,* local changes could still affect excitability and in turn structural remodeling (Butz et al., 2009). Finally, abnormal protein load could also affect local turnover by saturating the degradation machinery, which in turn could impair a remodeling process that is highly dependent on precise protein turnover (He et al., 2023). Together, our findings establish that the labeling strategy *per se* could affect structural plasticity and that avoiding interference with membrane composition is essential for preserving time of day dependent changes in structural plasticity.

Within this controlled methodological context, we next analyzed the influence of light and temperature on structural remodeling. Under the conditions examined, and contrary to our expectations, light intensity within the evaluated range did not significantly affect s-LNv terminal plasticity. Conversely, temperature exhibited a more pervasive effect, exerting its influence not on the remodelling process itself but on the mean complexity, with reduced temperatures giving rise to more complex terminals at both ZT2 and ZT14. The potential for both conditions to exceed the physiological range remains a possibility.

Finally, analysis of fine terminal structures provides further support for the hypothesis that the remodeling process is sensitive to experimental handling. The significant loss of filopodia following fixation and detergent-based permeabilization suggests that membrane protrusions are particularly fragile and that conventional immunolabeling protocols may underestimate their abundance. This methodological bias underscores the need for minimally invasive or live imaging approaches when studying structural plasticity at the subcellular level.

Overall, our results highlight that both environmental and methodological factors influence the interpretation of circadian structural plasticity. The combination of such experimental control with quantitative and unbiased tools such as *MorphoScope* will be essential to capture the mechanisms driving neuronal plasticity in adult circuits.

## Materials and methods

### Fly rearing

Male and female flies were maintained at 25°C on standard cornmeal yeast agar medium under 12:12 light-dark (LD) cycles, except for the experiments described in **Figure 3**. In these experiments, animals were reared at 22°C under low light intensity (70-280 lux) and LD 12:12 cycles. When adults were 3-5 days old, one group remained under those conditions, while others were transferred for 4 days to 22°C under high light intensity (1500-1600 lux), or to 25°C and 28°C under both low and high light intensities. Temperature treatments were performed in three separate incubators. Light intensity conditions were generated on different shelves within the same incubator by placing animals either on a shelf exposed to the incubator’s normal illumination or on another shelf where neutral density filters were applied to the incubator’s light tubes to reduce intensity (see the illustration on **Figure 3A**).

*Pdf*-RFP was provided by J. Blau (New York University, USA). *Pdf*-GAL4 (#6900, now available as part of #25031), *white*^1118^ (#5905) and UAS-CD8::GFP (#5137) were obtained from Bloomington Drosophila Stock Center (Indiana University Bloomington, USA).

### Immunofluorescence and imaging

#### Fixation protocols

To evaluate the effect of fixation on the preservation of the overall brain structure, we compared three protocols commonly used in the field. Flies were anesthetized with CO_2_ and decapitated and heads were then fixed in 4% paraformaldehyde (PFA, pH 7.5) either for 50 min at room temperature (RT) with gentle agitation, or kept on ice for 60 min in the same solution. Alternatively, whole bodies were briefly de-waxed in 96% ethanol and then fixed for 60 min in 4% PFA at RT with agitation. After fixation, brains were processed for immunofluorescence as described below and mounted in Vectashield (Vector Laboratories, USA). Images were acquired in a Zeiss LSM 710 (Zeiss, Germany).

#### Immunostaining and brain imaging

Male and female flies were anesthetized with CO_2_, decapitated, and their heads fixed for 50 min in 4% PFA at RT with agitation. Brains were then dissected in PBS containing 0.1% Triton X-100 (PT) and washed four times for 5 min. Samples were incubated for approximately 48 h at 4°C in PT 0.1% with 7% goat serum (Natocor, Argentina) and the corresponding primary antibodies (1:500). The primary antibodies used were chicken anti-GFP (Aves Labs, USA), rabbit anti-RFP (Rockland, USA), and rat anti-PDF (produced in-house, Depetris-Chauvin et al., 2011). After four additional 10 min washes in PT 0.1%, the samples were incubated for 2 h at RT with agitation with the secondary antibodies in PT 0.1% (1:500). Secondary antibodies used were Alexa Fluor 488 anti-chicken, Cy3 anti-rabbit, and Cy5 anti-rat (Jackson ImmunoResearch, USA). Brains were subsequently washed four times for 15 min in PT 0.1% and mounted in Fluoromount (Southern Biotech, USA) or Vectashield with the posterior surface oriented upward, using double-sided tape spacers (∼0.05-0.06 mm thick, applied in two layers; Rapifix, Argentina). The use of adequately thick spacers prevents tissue compression and preserves morphology. Mounting the brains with the posterior side facing the objective provides direct access to the s-LNv axonal terminals, which are located towards this surface, thereby reducing variability associated with imaging from different orientations. Approaching from the anterior side requires light to pass through a greater quantity of tissue, which increases fluorescence scattering and decreases image quality. Alternatively, samples can be mounted between two coverslips, thus enabling inversion of the preparation when required. See **Supplementary Figure 4** for details.

Confocal Z-stacks of the s-LNv dorsal terminals were acquired on a Zeiss LSM 710 using a 40X oil-immersion objective (NA 1.3) or on a Zeiss LSM 880 employing a 40X water-immersion objective (NA 1.2). Images were then used as the input for quantification of structural complexity using *MorphoScope* as described below. In **Figure 1**, full brain images are single snapshots, while the images of the terminals from all figures are the result of maximum intensity Z-projections of Z-stacks.

For comparisons between structural markers (**Figures 2A-2C**), laser power and gain were optimized for each brain to obtain the best GFP and/or RFP signals. Because PDF levels in s-LNv axonal terminals vary across the day, left hemispheres were imaged using the optimal settings for each brain, while right hemispheres were imaged using the parameters established from morning samples, when PDF expression is the highest. This approach allows assessing whether the use of an oscillating marker affects the quantification of structural complexity.

### Structural complexity quantification

The structural complexity of s-LNv axonal terminals was quantified using *MorphoScope*, an open-source graphical user interface (GUI) that implements and extends the three-dimensional spread quantification algorithm originally described by Petsakou et al., 2015. The software was developed in Python 3.9+ using *PySide6* for the graphical interface and *PyQtGraph* for real-time visualization, enabling intuitive operation by non-programmers while maintaining full computational transparency and reproducibility. Unlike the original MATLAB implementation which used JPEG-compressed images, our workflow processes uncompressed volumetric data (.czi, .lsm, .tiff) to preserve quantitative fluorescence information.

Upon loading a confocal Z-stack into *MorphoScope*, the user manually defines a region of interest (ROI) by drawing a polygon around the s-LNv axonal terminals, replicating the manual quality control step described in the original method. This step excludes unwanted structures and ensures that only the dorsal s-LNv arborizations are quantified. The selected volume can then be pre-processed using a combination of optional filters: Gaussian blur, median filtering, and intensity thresholding to reduce noise and background fluorescence. *MorphoScope* provides both manual threshold adjustment with real-time visual feedback and automated thresholding options (e.g., the Triangle algorithm), in contrast to the fixed 70% maximum intensity threshold used in the original MATLAB script.

Following Petsakou et al., 2015, the three-dimensional coordinate system is defined as follows: the Z-axis corresponds to the anterior-posterior axis as acquired by the confocal microscope (with the brain mounted posterior side up), while the X- and Y-axes are computed using principal component analysis (PCA) applied to the Z-projected fluorescence intensity. PCA identifies the direction of maximal arborization length as the X-axis, with the Y-axis defined orthogonally. This approach provides an objective, rotation-invariant coordinate system for each individual sample.

For each X-coordinate position, we calculated intensity-weighted mean Y and Z positions to define a 3D central curve through the arborization. Local spreads (standard deviations) in Y and Z were computed at each X-position, and global spreads (σ̄_x, σ̄_y, σ̄_z) were obtained by integrating these local spreads weighted by fluorescence intensity. The 2D spread (σ̄_x × σ̄_y) serves as the primary metric of structural complexity, as it captures the effective area of the axonal arbor while minimizing contributions from minor variations in Z-positioning across samples. Moreover, this measure is the most comparable to a Sholl analysis, which quantifies how terminals intersect successive concentric rings in the XY plane and does not account for the Z-axis.

The software calculates two complementary metrics to characterize the axonal arborization: (1) Integrated intensity (the sum of all fluorescence intensities across voxels in the thresholded 3D region), referred to as "axonal volume" by Petsakou et al. (2015), representing the total fluorophore content; (2) Geometric volume (number of voxels multiplied by voxel dimensions), representing the physical three-dimensional space occupied by the structure in µm^3^.

To enable co-expression analysis, *MorphoScope* includes a multi-channel quantification module. This allows for the simultaneous analysis of signal intensity in a secondary fluorescence channel (e.g., to quantify protein levels such as PDF) using the spatial coordinates defined by the primary morphology ROI. For these analyses, a maximum intensity Z-projection is generated from the secondary channel, and signal density is calculated as the mean intensity per occupied pixel (pixels with non-zero signal), reported in arbitrary units per pixel (AU/pixel) and per physical area (AU/µm^2^).

The complete source code, documentation, and standalone Windows executable are publicly available under the MIT license https://github.com/FranTassara/MorphoScope, (Tassara, 2025).

Terminal length was quantified in Fiji (Schindelin et al., 2012) by measuring the linear distance from the first branching point to the most distal tip of the terminal along the x-axis.

### Statistical analyses

The statistical analyses were performed in R software, version 4.5.1 (R core team, 2021) using the packages *tidyverse*, *nlme*, and *emmeans*, and ANOVA followed by t-test with Tukey corrections were performed. The Pearson correlation and Bland-Altman analyses were carried out using GraphPad Prism, version 8.0.2. Graphs were made in GraphPad Prism as well, and in all cases error bars indicate the standard error of the mean (SEM). Asterisks indicate statistically significant differences: * p < 0.05, ** p < 0.01, *** p < 0.001, and ns = not significant differences. Exact values for analyses and p values for contrasts are included in **Supplementary Tables 1-3**. The sample size (n) is indicated above the symbols in the graphs, and the type of sample is described in the corresponding figure legend.

Data were first inspected for normality and homogeneity of variances through graphical exploration. For data included in **Figure 2**, general linear models were fitted for each hemisphere and fluorescent marker to test for the effects of time point (ZT2 vs. ZT14), genotype, and their interaction on structural complexity. When visual inspection revealed heterogeneity of variance between time points, variance structures were modeled using the varIdent function to allow different residual variances per time point. The dataset shown in **Figure 3** was fitted with a general linear mixed model considering time point, light intensity, and temperature as fixed effects, and replicate as a random effect. To compare terminal metrics between mounting media (Fluoromount-G vs. Vectashield, **Figure 1**), an unpaired Student’s t-test was used for terminal length, whereas a Welch’s t-test was applied for complexity due to unequal variances.

To evaluate statistical power and effect size between time points, simulations were performed based on empirical data from five experimental replicates conducted at 25°C. Each experimental replicate included 10-29 animals. The spread of each brain was normalized to the mean of the experiment. In each simulation, n independent observations were extracted with replacement from each time point, and the means were compared using a t-test with Welch’s correction. This process was repeated 1,000 times for each sample size. Power was calculated as the proportion of simulations with p < 0.05, and the effect size was estimated using Cohen’s d for each repetition.

### *Ex vivo* preparations

Male and female flies were briefly anesthetized on ice, and brains were dissected in filtered medium containing 103 mM NaCl, 3 mM KCl, 5 mM TES, 8 mM trehalose, 10 mM glucose, 26 mM NaHCO_3_, 1 mM NaH_2_PO_4_, 4 mM MgCl_2_, and 1.5 mM CaCl_2_. Brains were mounted in the same medium following the procedure described above for fixed tissue and immediately imaged using a Zeiss LSM 880 with a 40X water-immersion objective. Representative terminals shown in **Figure 4A** are Z-projections of Z-stacks.

## Acknowledgements

We are grateful to the members of the Ceriani lab for their insightful discussions and to A. Liceri for providing fly food and assisting with fly work. We also thank A.H. Rossi and A. Ross from the Microscopy and Bioimaging Facility (Fundación Instituto Leloir) for their assistance and expertise in acquiring confocal microscopy images. MRC, JII, FJT and CCR are/were supported by graduate fellowships from the Argentine Research Council for Science and Technology (CONICET). FJT received a fellowship from the National Agency for the Promotion of Science, Technology and Innovation from Argentina (Agencia I+D+i). MFC is a member of CONICET. This work was supported by the Agencia I+D+i grant PICT2018-0995 and by NIH grant R01NS108934 (to MFC). The funders had no role in study design, data collection and analysis, decision to publish, or preparation of the manuscript.

## Supplementary information

**Supplementary Figure 1.**
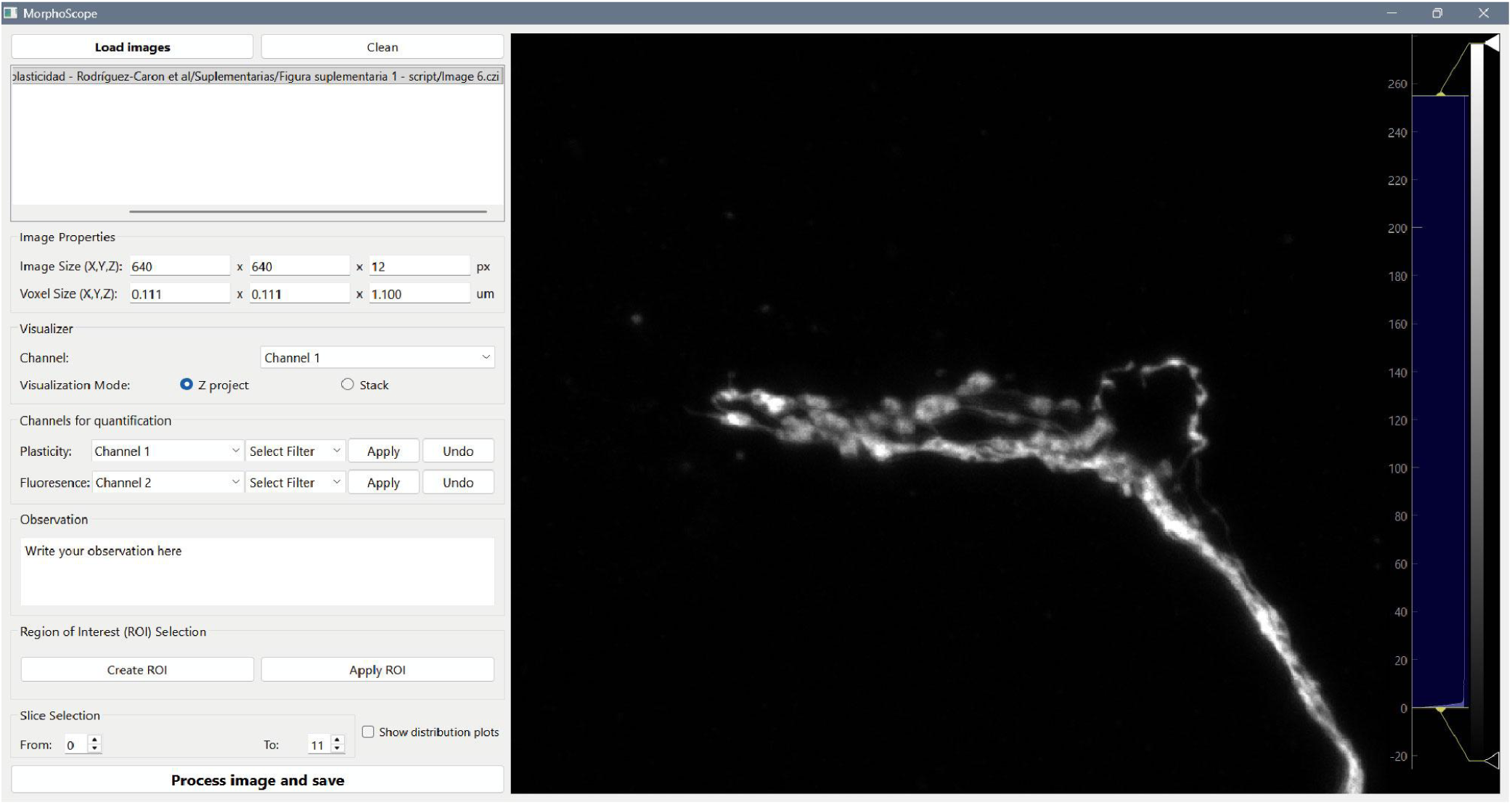
*MorphoScope* graphical user interface. The *MorphoScope* interface was developed to facilitate the complexity quantification of multiple terminal images based on the algorithm originally described in Petsakou et al. (2015). The interface is divided into functional modules, beginning on the left with “Load images” for selecting input files (.tiff, .czi, or .lsm), followed by “Image properties” which displays metadata like image and voxel sizes. The “Visualizer” controls the displayed channel on the right and selects between Z-projection or the full Z-stack visualization, while the “Channels for quantification” module specifies which channel is used for plasticity and which is used for fluorescence quantification, and also allows for filter application. Before processing, the user can define a note in “Observation”. Then, sets the area of interest using “ROI selection” and specifies the range of Z-planes via “Slice selection” to encompass the full terminal. Finally, quantification is executed by clicking “Process image and save” with results automatically added to an Excel file generated upon image loading. For further details, see https://github.com/FranTassara/MorphoScope.

**Supplementary Figure 2.**
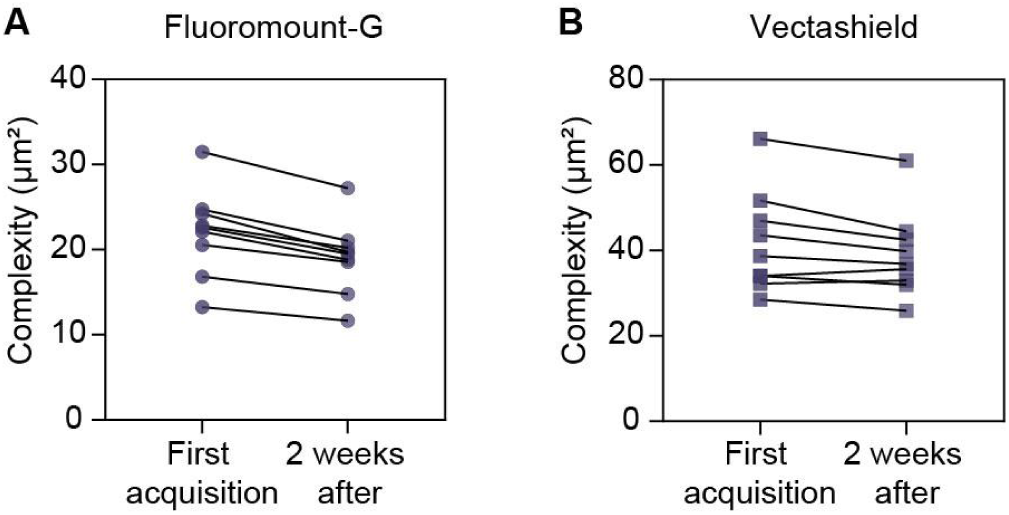
Stability of neuronal complexity over time in different mounting media. Comparison of the terminal complexity measured on the day after mounting (First Acquisition) and two weeks after for samples mounted in Fluoromount-G **(A)** and Vectashield **(B)**. Individual lines connect the same terminal re-acquired over time.

**Supplementary Figure 3.**
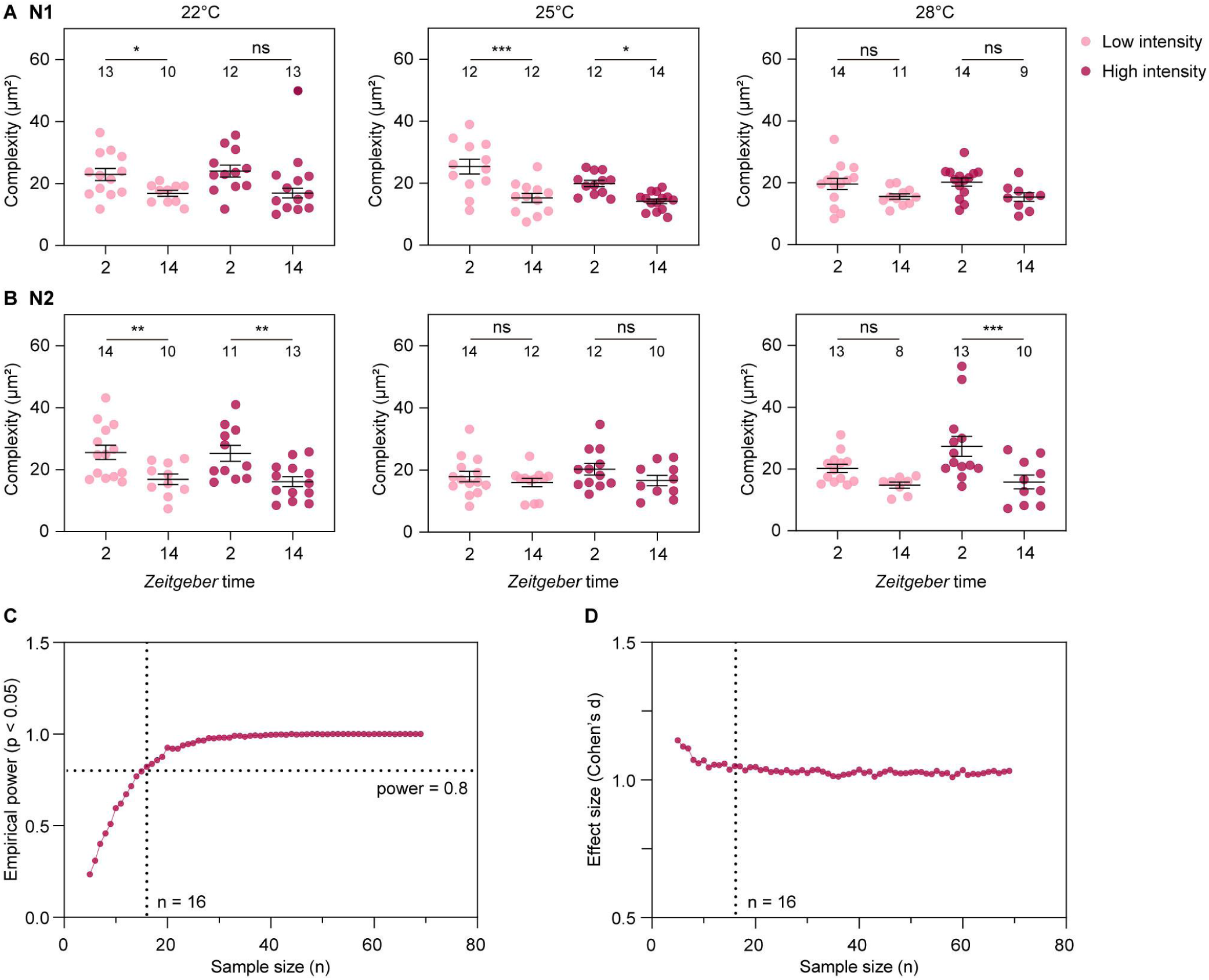
Sample size calculation. **(A and B)** Structural complexity quantification at different temperatures and light intensities. The panels display the two individual experiments that were shown pooled in Figure 3B. **(C)** Empirical power as a function of sample size. It shows the proportion of simulations yielding a statistically significant result (p < 0.05) across increasing sample sizes. The horizontal dotted line marks 80% power, and the vertical one indicates the sample size at which this threshold is achieved (n = 16). **(D)** Estimated effect size (Cohen’s d) as a function of sample size. It depicts the mean estimated effect size from repeated simulations across sample sizes. The vertical dotted line again marks n = 16 for reference.

**Supplementary Figure 4.**
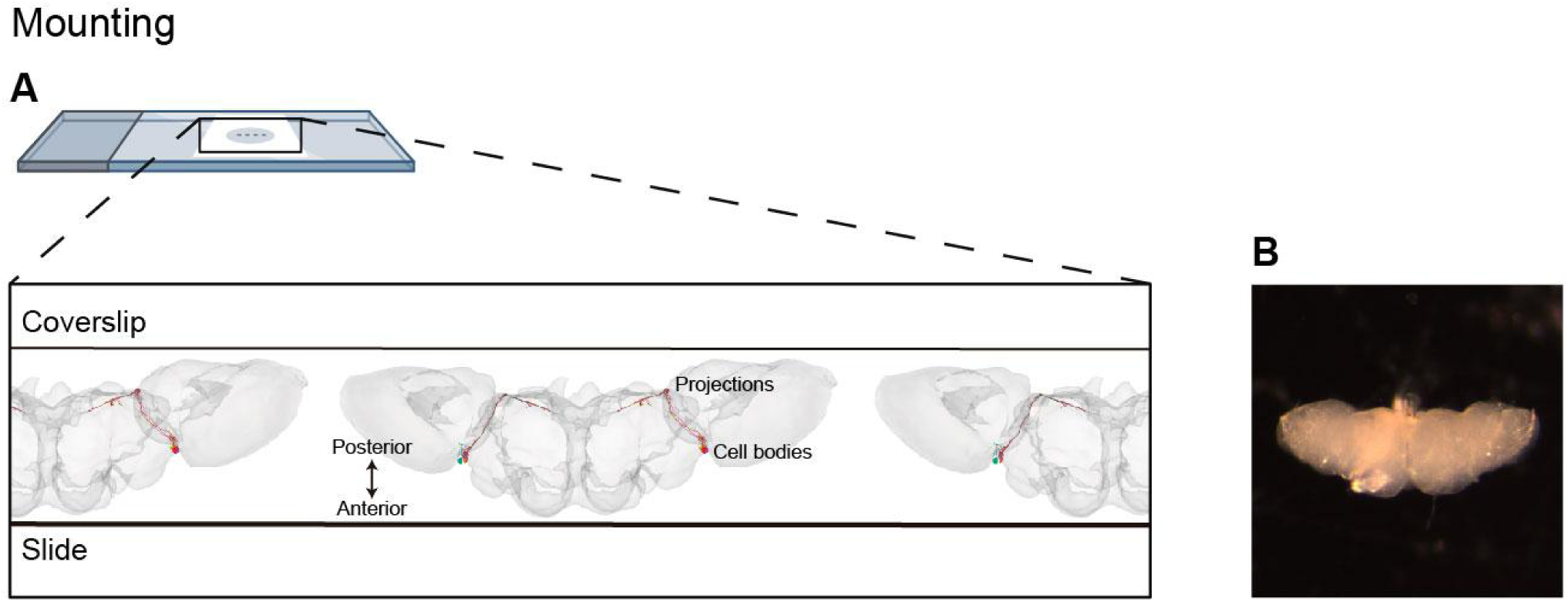
Schematic diagram of the brain mounting orientation. **(A)** A cross-sectional view illustrates brain placement between the glass slide and coverslip. Tissue is oriented along the anterior-posterior axis, with the anterior surface resting on the slide and the posterior surface facing the coverslip (and the microscope objective). The relative locations of somatic cell bodies and their neuronal projections are indicated to highlight the optical advantages of this mounting strategy. Alternatively, brains can be mounted between two coverslips to allow for inversion of the preparation depending on imaging requirements. Images of the 3D reconstructed brains were captured from the FlyWire Connectome Data Explorer (Matsliah et al., 2023; Dorkenwald et al., 2024). **(B)** Image of a real brain oriented as shown in **(A)**.

**Supplementary Table 1.**
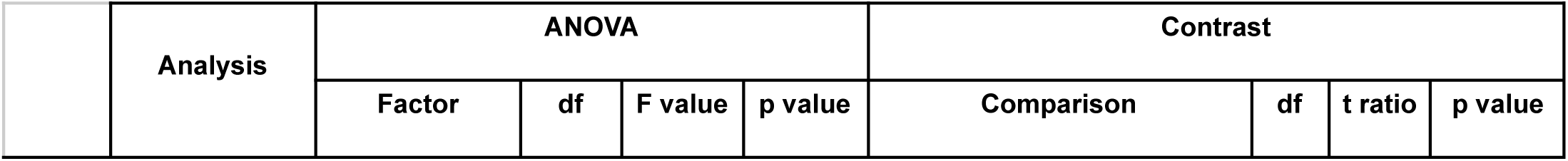
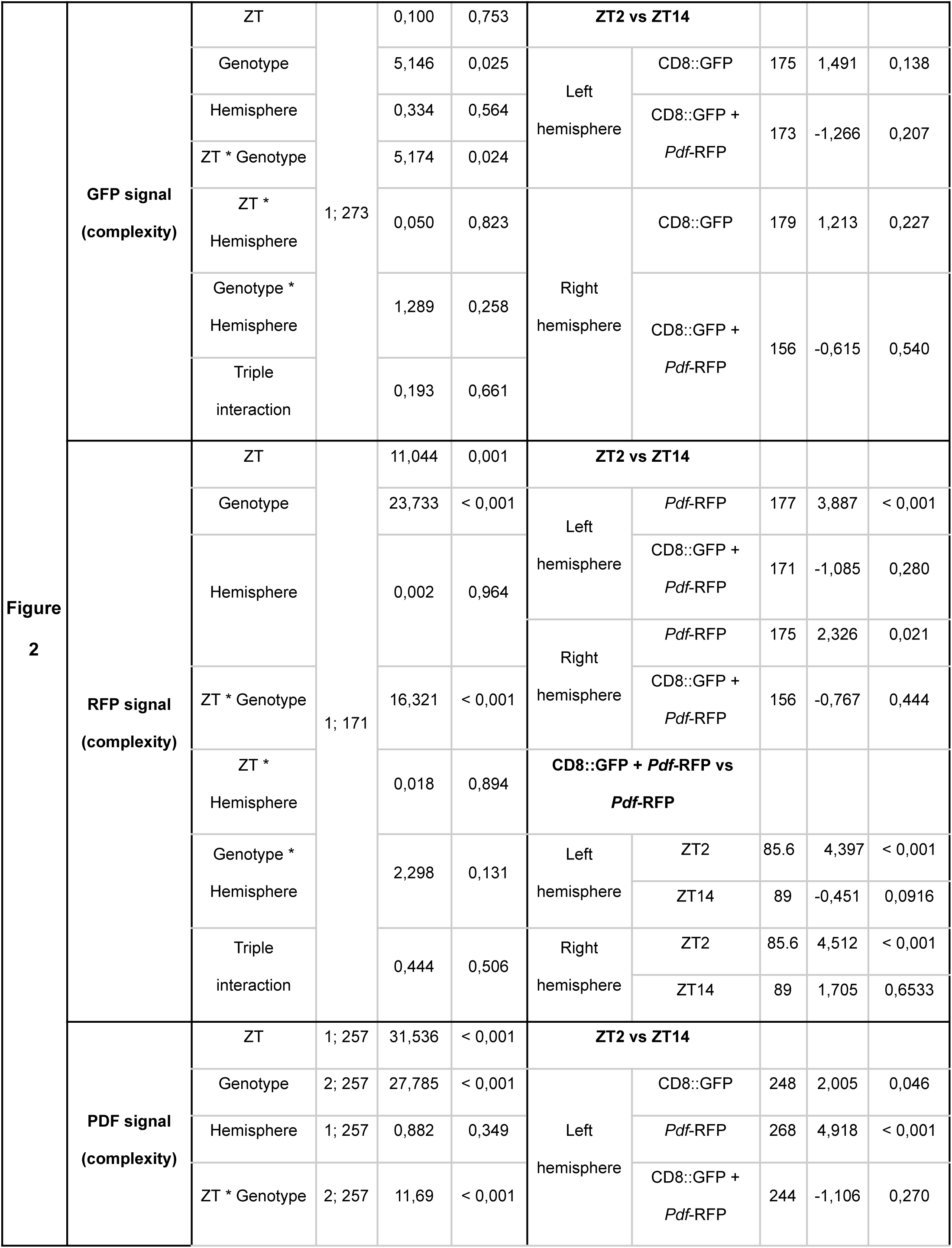
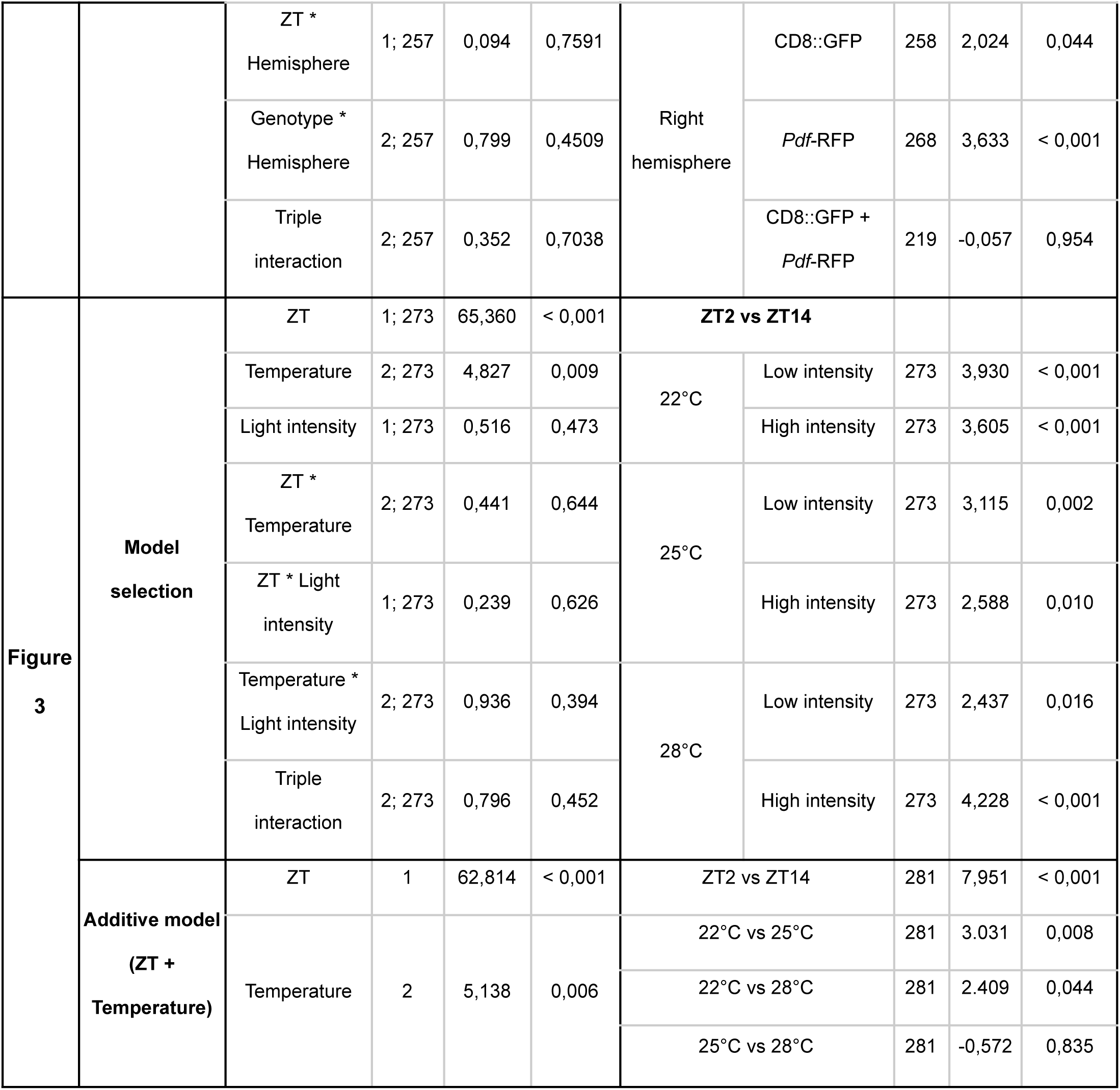
Detailed statistical outputs corresponding to the data presented in Figures 2 and 3. Summary of all general linear models, including main effects, interactions, and post-hoc contrasts for terminal complexity across different genotypes, times of day (ZT), and environmental conditions (temperature and light intensity). Test statistics (F values for ANOVAs and t ratios for contrasts), degrees of freedom (df), and exact p-values are provided for each comparison.

**Supplementary Table 2.**
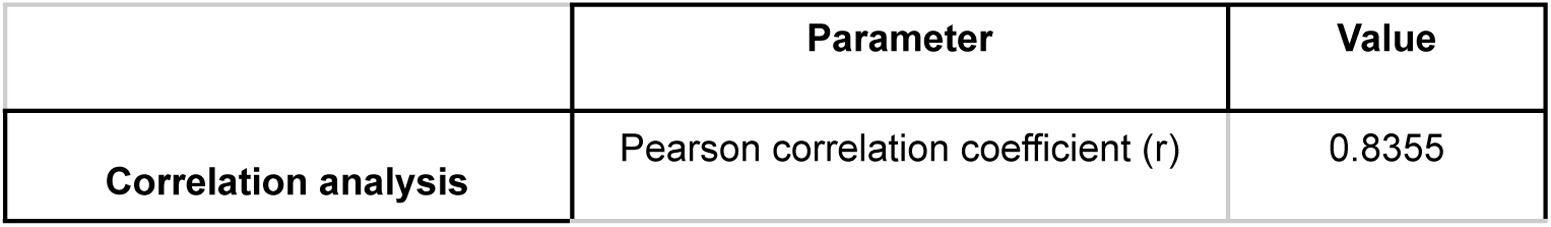
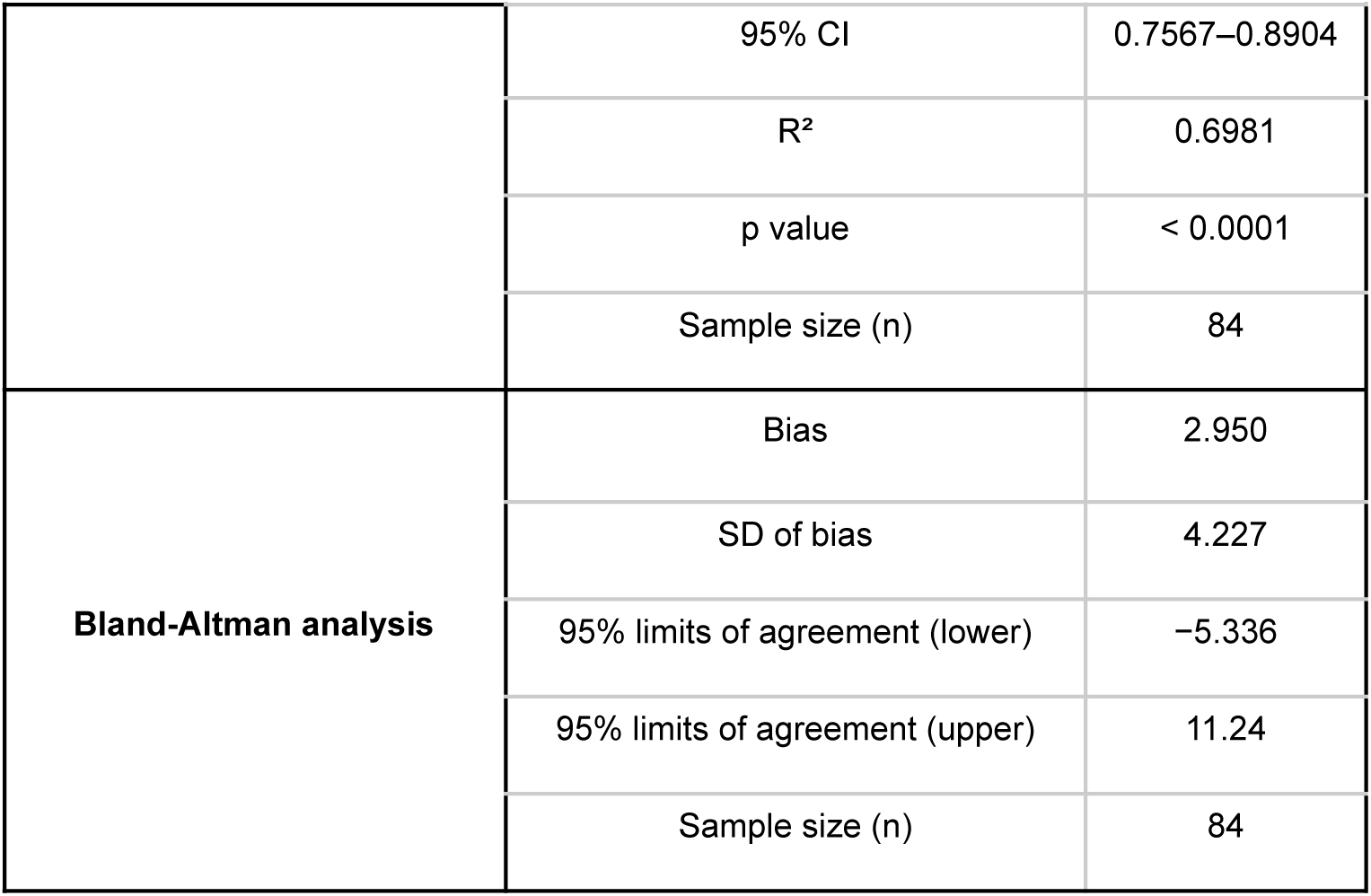
Method comparison using Pearson correlation and Bland-Altman analysis. Statistical parameters assessing the linear correlation (Pearson’s r, R^2^, and 95% CI) and absolute agreement (bias and 95% limits of agreement) between RFP and PDF signals for complexity quantification.

|  | Parameter | Value |
| --- | --- | --- |
| Correlation analysis | Pearson correlation coefficient (r) | 0.8355 |
|  | 95% CI | 0.7567–0.8904 |
|  | R<sup>2</sup> | 0.6981 |
|  | p value | < 0.0001 |
|  | Sample size (n) | 84 |
| **Bland-Altman analysis** | Bias | 2.950 |
|  | SD of bias | 4.227 |
|  | 95% limits of agreement (lower) | –5.336 |
|  | 95% limits of agreement (upper) | 11.24 |
|  | Sample size (n) | 84 |

**Supplementary Table 3.** Terminal metrics in Fluoromount-G vs. Vectashield mounting media. Values represent Mean ± SEM. Statistical differences were assessed via Welch’s t-test (complexity, XY spread) or unpaired Student’s t-test (terminal length).

| Variable | Fluoromount-G<br>(n = 22) | Vectashield (n<br>= 20) | Statistical Test | t | df | p-value |
| --- | --- | --- | --- | --- | --- | --- |
| **Terminal length** | 36.48 ± 1.359 | 49.18 ± 1.892 | Student's t-test | 5.53 | 40 | < 0.0001 |
| **Complexity** | 24.47 ± 1.679 | 44.93 ± 2.734 | Welch's t-test | 6.38 | 31.93 | < 0.0001 |

